# Identification of neuronatin as a SERCA2b regulin-like protein and assessment of its aggregation propensity via coarse grained simulations

**DOI:** 10.1101/2024.11.27.625357

**Authors:** Omar Ben Mariem, Lara Coppi, Emma De Fabiani, Ivano Eberini, Maurizio Crestani

**Author notes:** These authors equally contributed to this work (Omar Ben Mariem, Lara Coppi). These authors equally contributed to this work (Ivano Eberini, Maurizio Crestani).

## Abstract

Neuronatin (NNAT) is small transmembrane protein involved in a wide range of physiological processes, such as white adipose tissue browning and neuronal plasticity, as well as pathological ones, such as Lafora disease caused by the formation of NNAT aggregates. However, its 3D structure is unknown, and its mechanism of action is still unclear. In this study the two most well-known NNAT isoforms (α and β) were modelled and the interaction with the SERCA2b calcium pump was assessed using computational methods. First, molecular docking identified the same binding region as the one described for phospholamban, a thoroughly described SERCA inhibitor. Then, analyses of the flux of water molecules during molecular dynamics simulations highlighted significant similarities between the behavior of SERCA2b when in complex with phospholamban, and when in complex with either NNAT isoform. These results suggest that NNAT could be considered a “regulin-like” protein. Additional all-atom and coarse-grained simulations of multiple copies of NNAT highlighted a significant aggregation potential of both NNAT isoforms, supporting experimental data.

**Statement of significance:** This study presents the first structural model of neuronatin (NNAT) isoforms α and β. Through molecular docking and molecular dynamics simulations, we propose a NNAT interaction mechanism with the SERCA2b calcium pump similar to that of phospholamban, a known regulin and SERCA inhibitor. Our analyses also suggested a strong aggregation potential of NNAT based on all-atom and coarse-grained simulations, in line with experimental data on its involvement in Lafora disease. These insights suggest NNAT can be considered a “regulin-like” protein, advancing our understanding of its molecular function and contributing to new perspectives in targeting NNAT-related pathologies, as well as reinforcing the role of coarse-grained simulations as a valid tool to assess protein aggregation potential.

## Introduction

Neuronatin (NNAT) is a small transmembrane protein encoded in humans by the *NNAT* gene, whose structure is poorly understood. Although initially discovered to be mainly involved in brain development, over the years it has been shown to be involved in a wide range of physiological and pathological processes, e.g., white adipose tissue browning, glucose signalling, neuronal plasticity, and inflammation.(1) The *NNAT* gene is constituted by three exons and two introns, subject to alternative splicing, giving rise to a few isoforms, among which the isoforms α and β, constituted by 81 and 54 residues, respectively, are the most studied. Aberrant *NNAT* expression is correlated with several diseases, including diabetes, cancer, obesity, neurodegenerative disease, and retina degeneration.(2) Additionally, NNAT aggregates have been found to induce apoptosis, underlying diseases both in neuronal tissue (associated to Lafora disease), and in pancreatic β cells, leading to diabetes mellitus.(3, 4) Literature shows that both isoforms α and β can form aggregates in the cytoplasm without difference in propensity. Indeed, this phenomenon appears to be related to the increased expression levels of NNAT and/or proteasome dysfunction in pathophysiological conditions.(3, 5)

Choi and colleagues(6) recently demonstrated that cold exposure, which physiologically triggers thermogenesis, reduces the expression of NNAT in white adipose tissue (WAT) through ubiquitination and subsequent degradation of the protein. Furthermore, genetic inactivation of the *Nnat* gene in mice promotes browning of subcutaneous WAT stimulating the uncoupling protein 1 (UCP1)-independent thermogenesis.(7, 8) This alternative type of thermogenesis is distinguished from the UCP1-dependent thermogenesis because the electron transport chain (ETC) is coupled with oxidative phosphorylation. The ATP produced is then used to fuel a futile cycle of calcium ion (Ca^2+^) transport between the endoplasmic reticulum (ER) and the cytoplasm under the action of the sarco-/endoplasmic reticulum Ca^2+^-ATPase 2b pump (SERCA2b) and the ryanodine receptor 2 (RyR2).(6, 9)

SERCA2b is a member of the SERCA pump family, P-type ATPases transporting Ca^2+^ from the cytoplasm to the ER Three genes, i.e., SERCA1, SERCA2, and SERCA3 are responsible for the production of the 11 isoforms, which share the same function but differ in tissue distribution.(10) They have a fundamental role in regulating Ca^2+^ compartmentalization, necessary for a wide array of physiological processes in all tissues.(11–13) Therefore, the regulation of SERCA activity profoundly impacts cell homeostasis. Phospholamban (PLB) and sarcolipin are two of the key regulators of the activity of SERCA pumps, called “regulins”.(14) These small proteins are present in skeletal muscle fibres and cardiomyocytes, where they inhibit the activity of the various SERCA isoforms, thereby increasing the cytosolic concentration of Ca^2+^ ions and, among other effects, promoting contractions of these fibres.(15–27)

Like all pumps, SERCA works by undergoing a series of consecutive conformational changes that results in the ATP hydrolysis and Ca^2+^ transport. It has been proposed that PLB and sarcolipin stabilize one of such conformations, thereby preventing the correct pump function. (2, 6, 28) It has been recently shown that PLB undergoes phosphorylation at serine residues that negatively affects its inhibitory activity on SERCA2A.(23)

NNAT is thought to have a role comparable to the ER-membrane protein PLB, decreasing the transport rate of Ca^2+^ from the cytoplasm to the ER. Interestingly, a recent phosphoproteomic study identified a phosphorylation site on Ser56 of the α isoform, corresponding to Ser29 of the β isoform, which might also suggest a regulatory mechanism similar to that of PLB.(29)

In this context, this study aims at proposing a potential interaction mechanism between NNAT and SERCA2b, to characterize the inhibition of the Ca^2+^ pump, which could become an interesting potential target for drug discovery. Because of the lack of experimentally solved structures of either NNAT α or β, the first step of the study was the generation of a 3D model for both isoforms. After obtaining a reliable conformation of these structures, it was possible to assess the potential inhibition mechanism, comparing it to the one described for PLB. Additionally, these models were used to study the formation of protein aggregates, involved in neurological and metabolic diseases.

## Material and Methods

### Modeling of human NNAT, SERCA2b, and PLB

Because no experimental structures for human NNAT or SERCA2b were available, modeling procedures were used to obtain reliable 3D structures through the following steps. The sequences of the two human NNAT isoforms (UniProt ID: Q16517) were retrieved from the Uniprot knowledgebase database.(30) A protein BLAST(31) search in the Protein Data Bank (PDB)(32) for a NNAT homolog to be used as a template for homology modeling returned no results for either isoform.

Preliminary secondary structure predictions were run using TMHMM(33) and PsiPred.(34) AI-based *ab initio* strategies were subsequently employed to obtain a reliable model, specifically, RosettaFold(35) and AlphaFold2.(36)

The BLAST search performed using the sequence of human SERCA2b (UniProt ID: P16615) returned numerous results. Among the available structures that could be used as templates, the *O. cuniculus* SERCA1a structure was selected (PDB code: 4KYT(37)), with an identity percentage of approx. 80% and similarity of approx. 89%, making homology modelling the most appropriate modeling procedure. This specific structure was chosen because it is already in a complex with PLB, making the following comparative steps more reliable. The human PLB model (UniProt ID: P26678) was also built via homology modelling by first superposing the transmembrane helix of the full PLB monomer from *O. cuniculus* (1N7L(38)) on the PLB structure available in the 4KYT complex, then using that structure as template, with an identity and similarity of approx. 91%.

### Equilibration and Cluster Extrapolation

Molecular dynamics (MD) simulations and frame clustering procedures were carried out with Desmond and the Schrödinger Small-Molecule Drug Discovery Suite 2022-3 (D. E. Shaw Research, New York, NY; Schrödinger, New York, NY).(39)

The Desmond System Builder tool was used to place the two obtained NNAT models into a 1-palmitoyl-2-oleoyl-sn-glycero-3-phosphocholine (POPC) membrane bilayer. Protein orientation was set up according to the OPM server, which provides spatial arrangements of membrane proteins with respect to the hydrocarbon core of the lipid bilayer. The system was solvated with SPC water molecules in an appropriately sized orthogonal box (a buffer of 15Å in the x, y, and z dimensions). Sodium and chloride ions were added to reach a 0.15 M concentration and neutralize the system. The system was energy-minimized to relax the assembly and remove clashes between protein, membrane, and solvent in the new setup.

One 1000 ns MD simulation was run for each isoform to obtain equilibrated and more realistic structures. Periodic boundary conditions (PBCs) and the following parameters were set: 300 K and Nose-Hoover thermostat for temperature coupling, 1 bar and Martyna-Tobias-Klein piston for pressure coupling, and 2 fs as the integration time step. Coordinates and velocities of each atom were saved every 0.5 ps. The OPLS4 force field was used to parametrize the atoms for the MD simulations.(40)

One additional MD simulation for each isoform was run keeping the protein in solution, to obtain soluble conformations that could be used for the following MD simulations to assess their aggregation potential.

Root mean square deviation (RMSD) and solvent accessible surface area (SASA) were calculated using the Schrödinger Python API. RMSD distances between MD frames were used to perform clustering and obtain a representative structure for both the isoforms.

### Aggregability assessment with all-atom and coarse-grained MD simulations

The most representative medoid of the most populated cluster obtained from the MD simulations run in solution was obtained for each isoform. Two systems were built inserting seven copies of each one in a solvated box. For all-atom MD simulations, 400 ns were run using the previously described parameters.

To obtain the coarse-grained systems, the SIRAH conversion tool available with the AmberTools22 suite was used to convert the groups of atoms in the structures to coarse-grained beads. The SIRAH force field was used to parametrize the systems. MD simulations were run for 1000 ns using the pmemd engine available in the Amber22 suite using Langevin dynamics. (41–43)

### Complexes Generation and Analysis

The medoid of the most representative cluster for each isoform was extracted from the in-membrane MD simulations and used as starting point for protein∷protein docking calculations using Piper. (44) The procedure was performed using the default Piper parameters, obtaining 10 final poses. No restraints to narrow down the possible interaction interfaces were defined. Because of the high flexibility of the C-terminal portion of both isoforms, only the identical transmembrane helix was considered.

The full NNAT isoforms were then rebuilt, minimizing the complex to make sure to avoid clashes. The proteins were then inserted in a POPC bilayer and three replicas of 500 ns MD simulations were run as previously described.

The analysis of the interactions between the two proteins and the calculations of the fluxes of water molecules were performed using the Schrödinger Python API.

## Results and discussion

### Model generation and equilibration

As previously mentioned, there seems to be consensus regarding the overall architecture of the two NNAT isoforms.(3) Specifically, both are characterized by a transmembrane helix, an unstructured portion of different length in the two isoforms, and a terminal cytoplasmic helix. Preliminary predictions using TMHMM and PsiPred support these hypotheses. (Fig. S1) Since homology modeling was not possible due to the lack of appropriate templates, *ab initio* methods were applied using AI-based software, specifically RosettaFold and AlphaFold2 (Fig. 1). While displaying significant similarities, AlphaFold predicted an unrealistic conformation for the α isoform (panel A, left side), with the hypothesized cytoplasmic α-helix expressed by the third exon parallel to the N-terminal, which would therefore also be embedded in the membrane. Additionally, the α isoform model obtained by AlphaFold was the only one presenting outlier backbone dihedral angles values in the Ramachandran plot (Fig. S2). Because of this, to avoid introducing unnecessary bias in the following steps, the models obtained by RosettaFold were selected. Interestingly, the overall predicted architecture strongly resembles the one of PLB and sarcolipin, although in those proteins the transmembrane helix is at the C-terminal.

**Figure 1.**
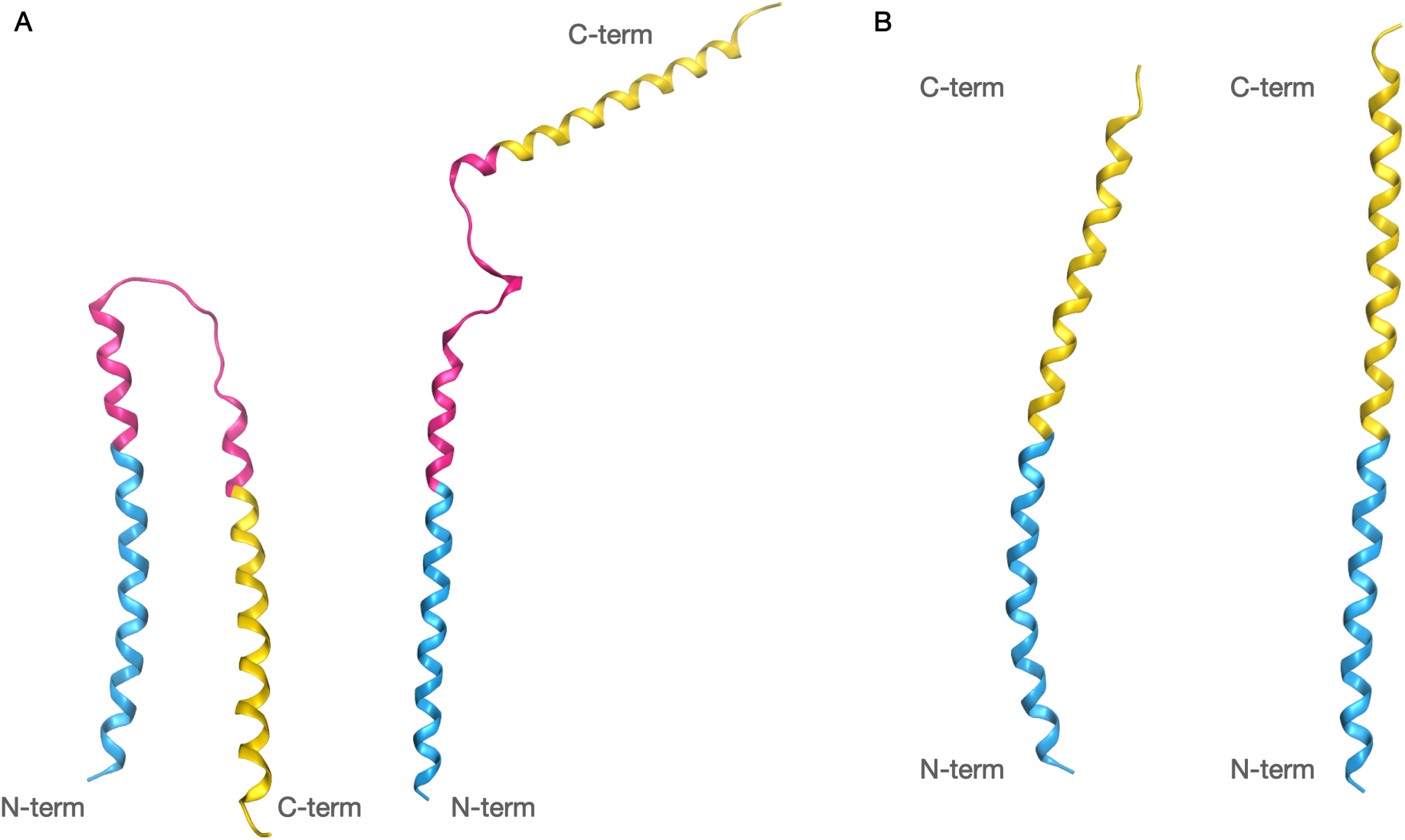
Structure predictions of NNAT α (A) and NNAT β (B) with AlphaFold on the left and RoseTTAFold on the right of each panel. While for the β isoform the two software demonstrate a very similar prediction, the models of the α isoform present very different conformations. However, interestingly, the two algorithms provide similar predictions of both the helical and unfolded portions. All structures are coloured to reflect the exons in the NNAT DNA (cyan for exon 1, magenta for exon 2, absent in the β isoform, and yellow for exon 3).

As the models could not be considered reliable without further investigations, MD simulations were run to better reflect the physiological conditions and allow the model to relax and find a more stable conformation. Figure 2 reports the RMSD profiles of the proteins during the simulations. It is possible to observe how both isoforms, after an initial significant structural rearrangement (approx. 150 ns), which was expected due to the highly mobile structure, reached a plateau which was maintained throughout the simulations.

**Figure 2.**
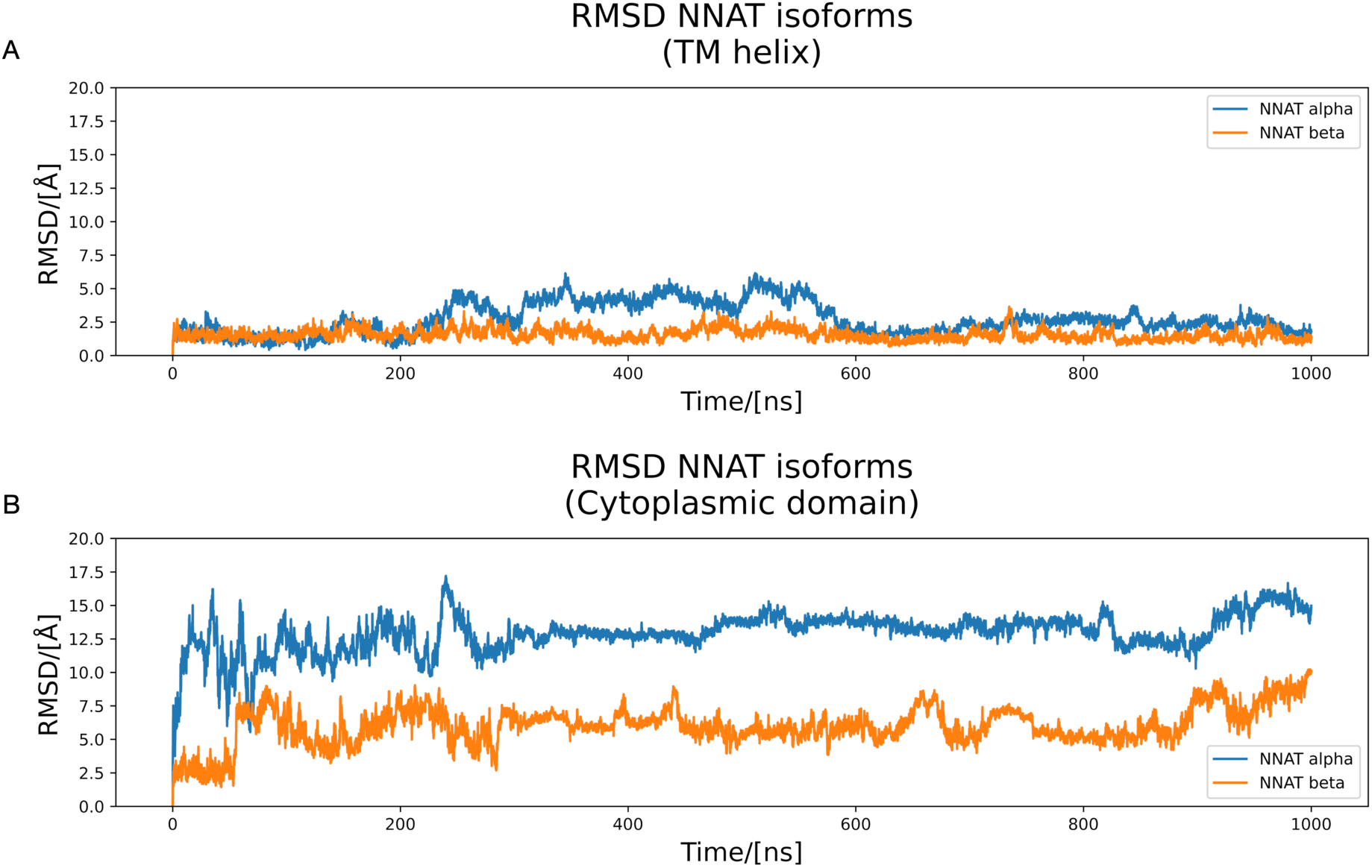
RMSD profiles of the two NNAT isoforms during the MD simulations in the membrane. A) RMSD of the transmembrane (TM) α-helices. The profile appears flat with relatively low values, confirming stability of the models in that portion. B) RMSD of the cytoplasmic domain after alignment with the transmembrane α-helices. The values of RMSD are high because of the significant relative mobility between the cytoplasmic and transmembrane domain, but the profiles appear flat, demonstrating good overall stability.

The secondary structure analysis confirmed that the largest portion of both isoforms retain the hypothesised α-helical structure, although the large loop in the α isoform, as well as the first 10 residues in the β isoforms are indeed in a random coil conformation (Fig. 3). Overall, these simulations indicated that the two isoforms, after an initial structural rearrangement, were stable and maintained a high α helical secondary structure propensity.

**Figure 3.**
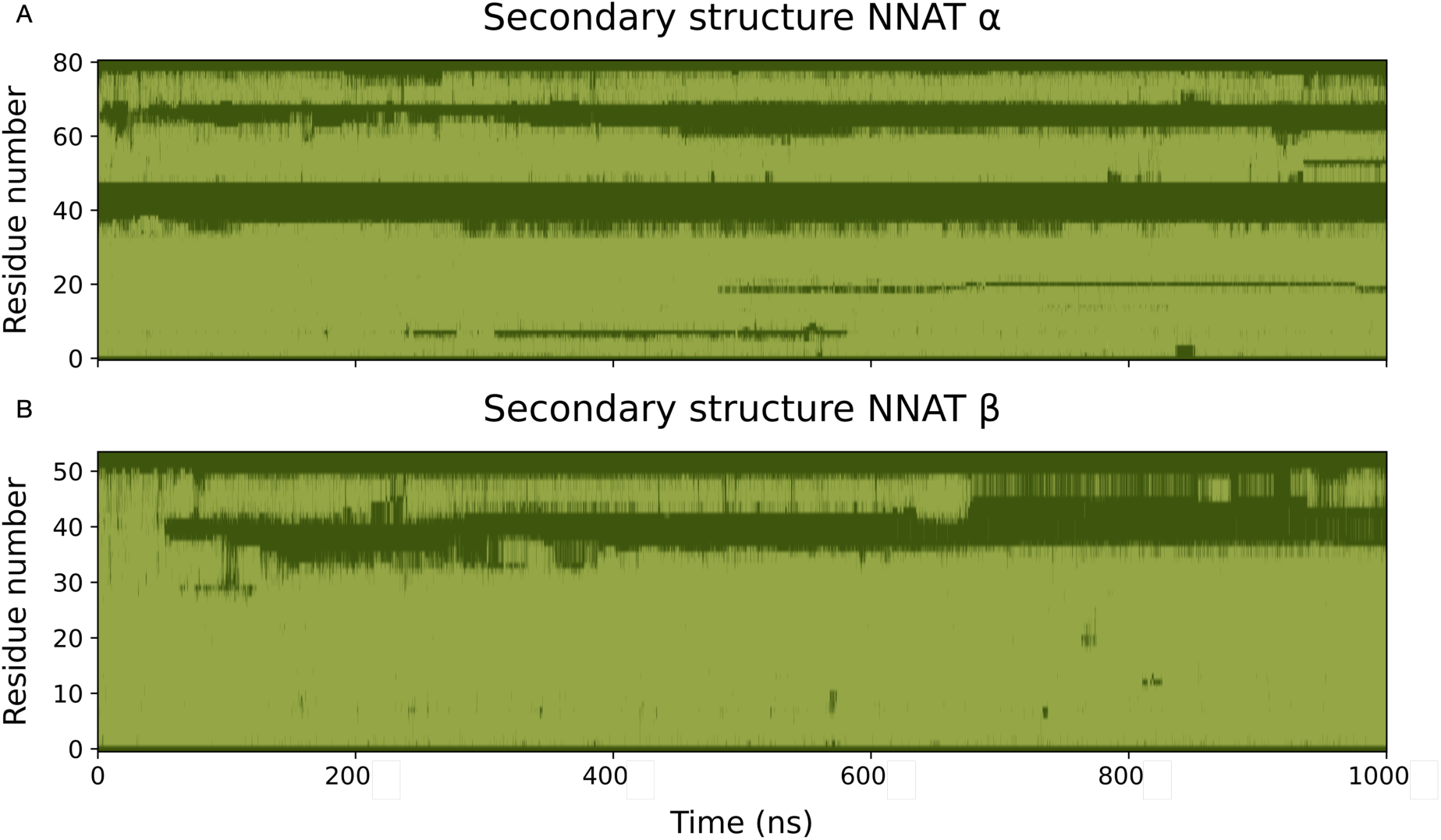
Secondary structure of the two NNAT isoforms. Light green indicates α-helical secondary structures, while dark green indicates an unfolded portion of the protein. A) Secondary structure of NNAT α. The model retained a stable transmembrane α helix for the whole simulation up to residue 35. The cytoplasmic portion alternates unfolded portions to α-helical ones. B) Secondary structure of NNAT β. The model retained a stable α helix for most of the simulation up to residue 35. Despite the starting conformation as a fully helical structure, most of the cytoplasmic portion loses its secondary structure during the simulation.

To evaluate the mechanism of the inhibition of SERCA2b by NNAT, the calcium pump model was also generated. In this case, a homology modelling procedure could be applied, using a template with an extremely similar sequence (4KYT, see Fig. 4 and alignment in Fig. S3). Because the template is already in a holo-conformation with PLB, the obtained model was simply energy minimized and no further MD simulations were run on the model to be used for the protein∷protein docking, to maintain the most suitable conformation for protein binding and to avoid bias introduction. SERCA2b is the only SERCA isoform with 12 transmembrane α-helices but, since the template does not have the extra helix, that helix was not modelled. This was done to avoid biases due to the unreliable conformation of the modelled structure. The absence of the helix is not a concern, as the mechanism of action of proteins regulating SERCA function is shared among all isoforms, and the binding site is far from the hypothesized position of the extra α-helix. A MD simulation of the model was then run to assess the water molecule fluxes in the pump, which will be discussed in a following section.

**Figure 4.**
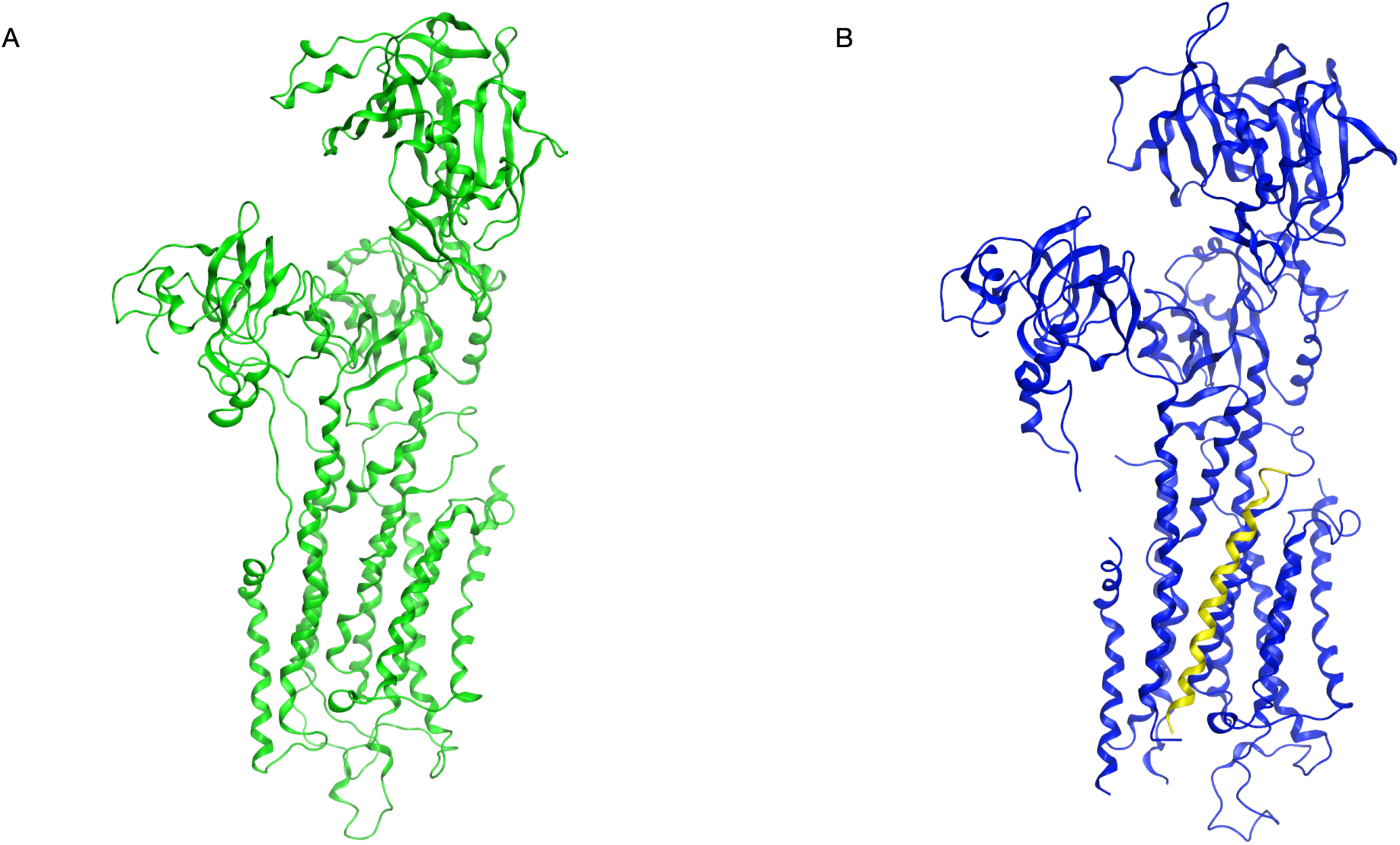
(A) 3D model of human SERCA2b and (B) crystal structure (PDB code 4KYT) of the *O. cuniculus* SERCA1b (blue) co-crystalized with its inhibitor PLB (yellow).

The same structure (4KYT) was used to build a complete human PLB model. Although an experimentally solved structure of human PLB is already available, it is only present in its pentameric form. Therefore, first the monomeric PLB experimental structure of *O. cuniculus* (1N7L) was superposed to the truncated structure present in the 4KYT crystal, then the human model was generated via homology modeling. Because of the high sequence identity of both proteins, the SERCA2b∷PLB humanized complex can be considered reliable. A MD simulation was further run to assess the flux of water molecules in SERCA2b, which will be discussed in a later section.

### NNAT isoforms aggregation

Aside from its numerous physiological roles, NNAT has also been found to cause significant cell apoptosis and tissue degeneration in both brain and pancreas due to the formation of toxic aggregates. To assess the aggregation potential of the two NNAT isoforms, MD simulations were run. Initially, a 1 ns long MD simulation of the two models were run in solution, to obtain conformations resembling those adopted by the proteins in the cytosolic compartment, for example, due to overexpression and/or impairment in their degradation pathway. Expectedly, as they were not in their physiological environment, across the membrane, both NNAT isoforms lost some of their secondary structure features and reached a more compact conformation, as seen from the SASA and the radius of gyration (Fig. 5). This is especially evident for the α isoform, while the β NNAT seems to keep a fairly stable level of compactness with small fluctuations over the whole simulation. These significant rearrangements can be seen especially in the first 10 residues of the N-terminal portion for both isoforms, where it is possible to observe a significant loss of secondary structure, as the hydrophobic domain, physiologically inserted in the phospholipid bilayer, unfolded. The conformations of the final frames were used to build systems containing 7 copies of each NNAT isoform, randomly placed in a fully solvated box.

**Figure 5.**
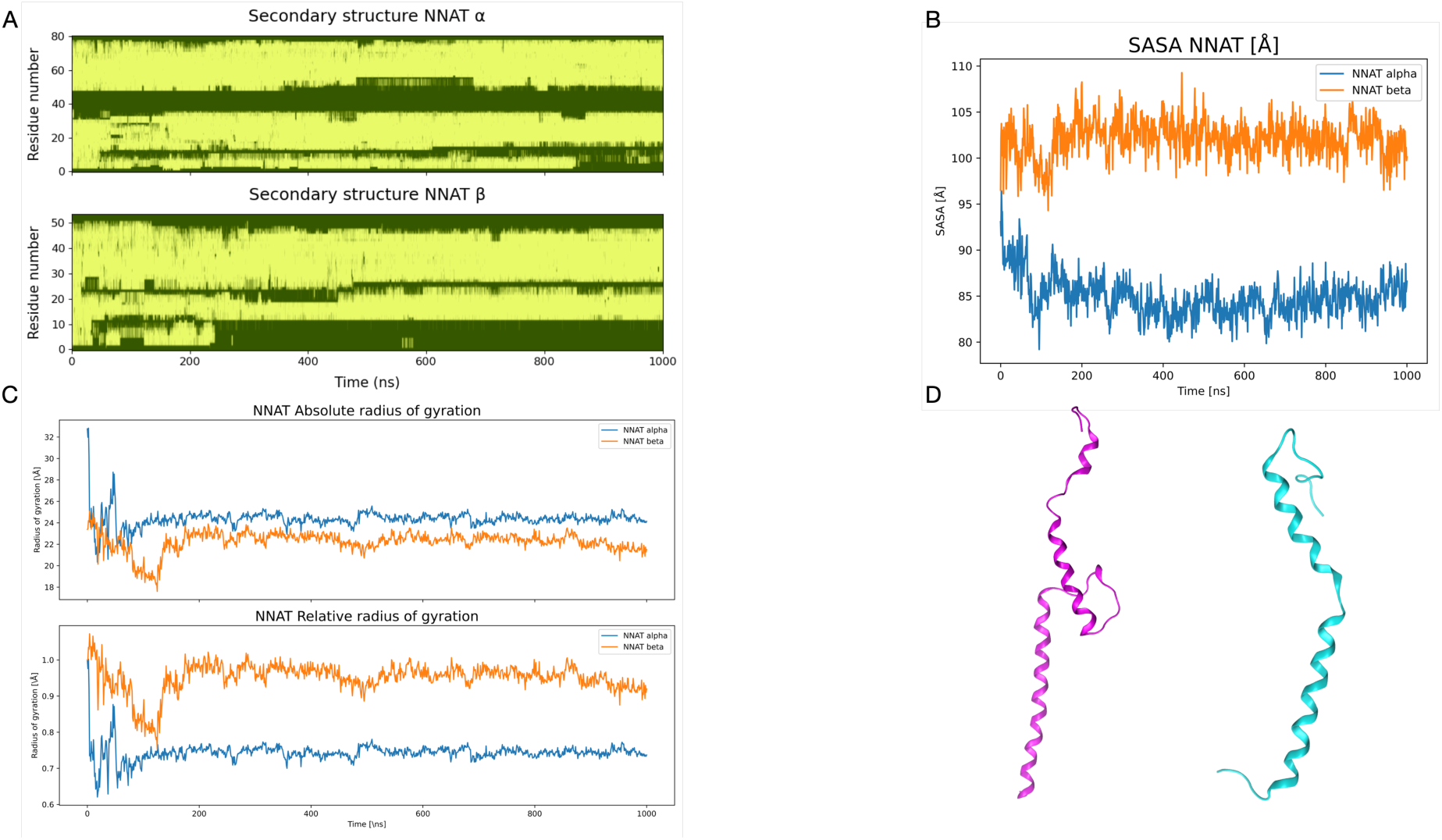
Analysis of the MD simulations of the two NNAT isoforms in solution. A) Both isoforms tend to lose some secondary structure features, especially at the N-terminal, which is hydrophobic and should be transmembrane. B) SASA of the two isoforms. C) Radius of gyration of the two isoforms, both in absolute and relative values (compared to the first frame). The decrease in radius of gyration is evident for both simulations, but especially for the α isoform, due to its higher degrees of freedom. D) Final structure of the two isoforms (purple: α, cyan: β). For both structures, the N-terminal is at the bottom of the figure.

From the same starting coordinates, two systems were generated: an all-atom system and a coarse-grained one using the SIRAH tool and force field. This choice was made to compare the results obtained by different methods, which could further demonstrate the potential aggregation of these proteins. The results for the all-atom simulations shown in Fig. 6 highlight that aggregation does indeed happen in a short time, as at around 300 ns one unique aggregate can be observed for both NNAT isoforms, and it is maintained for the duration of the simulations.

**Figure 6.**
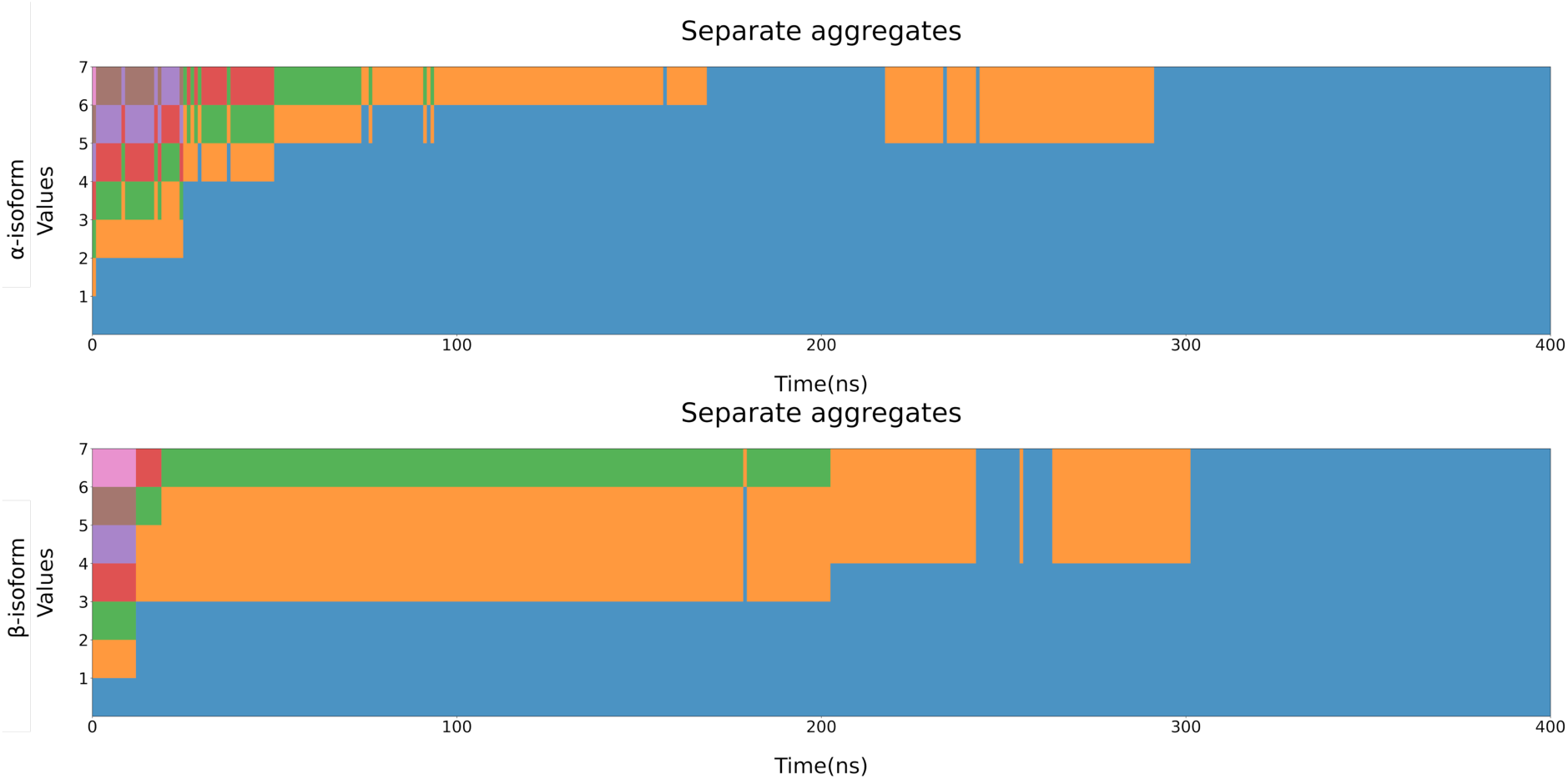
Separate entities observed for the all-atom simulations. Each colour represents a unique “entity”, defined as a copy or ensemble of copies not interacting with the others. It is possible to see that, for both the isoforms, copies begin to aggregate rather quickly and after 300 ns all copies are already fully aggregated and remain as such for the rest of the simulation.

Interestingly, in the coarse-grained simulations, even after 4 μs the copies are not fully aggregated, as only smaller aggregates are formed (Fig. 7). This could be due to the approximation introduced by the coarse-graining process, or by the higher friction coefficient used for these simulations. This process could be further studied using additional simulations with different coefficients and different NNAT concentrations, however the all-atom simulations appear to confirm a high aggregation potential for both isoforms, supporting the already available experimental evidence.(3)

**Figure 7.**
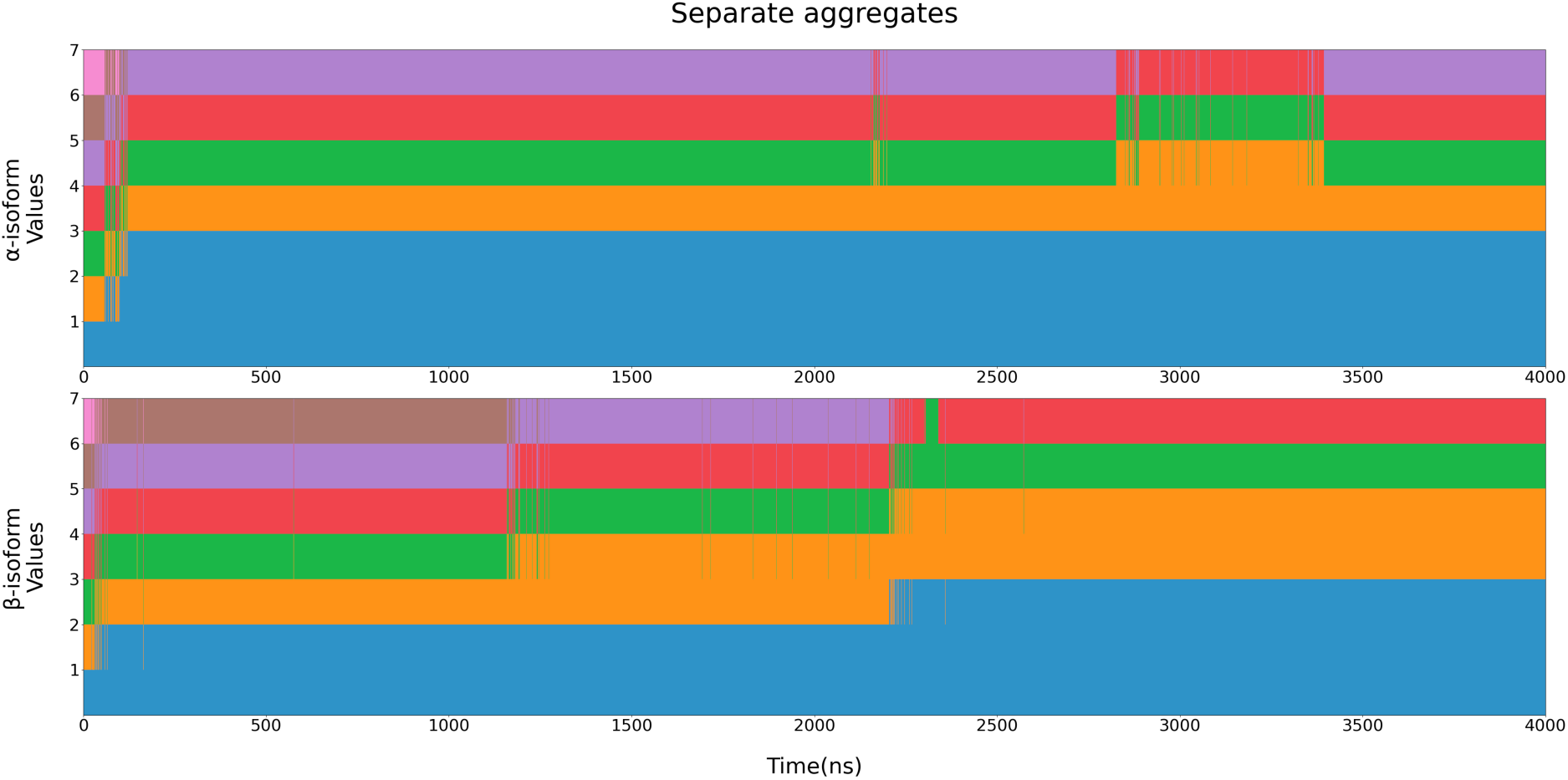
Separate entities observed for the coarse-grained simulations. Each colour represents a unique “entity”, defined as a copy or ensemble of copies not interacting with the others. Differently from the all-atom simulations, it is not possible to observe complete aggregation, even after 4 μs.

### Identification of the SERCA2b∷NNAT binding site

After extracting structures of both isoforms from membrane MD simulations, initial attempts of performing protein∷protein docking to obtain SERCA2b∷NNAT complexes were performed. Expectedly, because of the bulky cytoplasmic terminal of both isoforms, and the overall static nature of this procedure, no poses were found. Therefore, to avoid biases caused by the specific frame chosen to extract the structure, only the transmembrane helix (expressed by the first intron, identical in both NNAT isoforms) was used for the calculations. By doing this, several extremely similar poses were obtained, each representing a differently sized cluster of poses (Table 1). Considering the 8 most populated clusters (>95% of the total poses), all are located in the groove between helices M2, M6, and M9, extensively characterized as the PLB binding site (Fig. 8). (15, 22, 23, 37)

**Figure 8.**
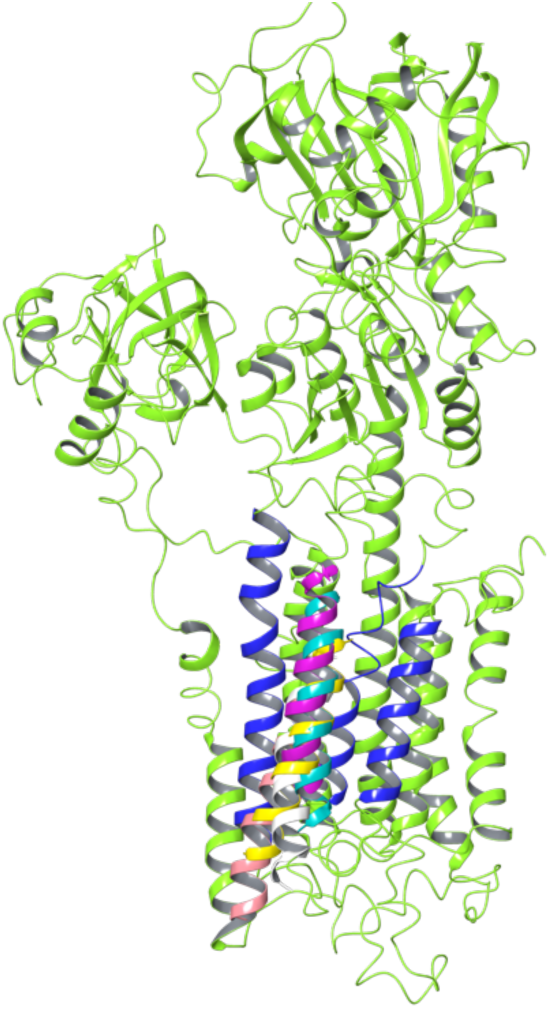
(A) Top ranking protein::protein (PIPER) docking poses. All the represented poses are found in the well characterized M2-M6-M9 groove (colored in the darker blue hue).

Interestingly, the highest scoring poses were characterized by having the NNAT helix oriented in the opposite direction of PLB, which is with its N-terminal towards to lumen and the C-terminal towards the cytoplasm. This also supports hypotheses made during the years by researchers, pointing towards NNAT having the C-terminal tail in the cytoplasm, potentially able to interact with the ATP-ase and translocation domains of SERCA and having, overall, a similar effect on SERCA activity to the more well-known and characterized regulins.(45) The way this result obtained by a fully unbiased molecular docking procedure correlates with experimental data supports the reliability of the hypothesized binding site and the orientation of the transmembrane helix.

### SERCA2b water fluxes regulation

The two NNAT isoforms were then “rebuilt” starting from the highest scoring docking pose superposing the selected representative structures to the docked transmembrane helix. Because of numerous clashes between the C-terminal of NNAT and the cytoplasmic domains of SERCA2b, that specific portion was energy minimized to better fit in that specific conformation (Fig. S2). These two complexes, alongside the apo-SERCA2b and the SERCA2b∷PLB complex, were inserted in a POPC bilayer and solvated. Three replicas of MD simulations for each system were performed. The data obtained by the apo-SERCA2b system could be considered as a “negative control”, since it simulates how the protein would behave in physiological conditions. On the other hand, the SERCA2b∷PLB complex could be considered as the “positive control”, since there is wide experimental evidence regarding its effect on the Ca^2+^ pump.

During these MD simulations, the most noticeable difference between the complexes was the regulation of the water pathways and, by proxy, of the Ca^2+^ ions. In fact, literature suggests the presence of two cytoplasmic entries (an “N” and a “C” path), and one endoplasmic exit path.(46) The simulated complexes show different patterns of the opening of such paths, suggesting a potential effect of regulins on SERCA2b function through this mechanism.

Fig. 9 represents the water fluxes through the two cytoplasmic paths and the endoplasmic exit. Interestingly, the apo-SERCA2b behaves as expected, showing a direct connection between the cytoplasm and the ER. Differently, during the simulations of all complexes with PLB and the two NNAT isoforms, this connection is almost severed, with the secondary cytoplasmic opening forming, giving water molecules a way back into the cytoplasm. This can be considered a mechanistic explanation on the role of regulins on SERCA function. This behaviour is especially notable in the PLB system, which is known to be the strongest inhibitor of the Ca^2+^ transport. A difference in magnitude of the effect appears to be visible between the two NNAT isoforms, with the α one causing a marginally larger secondary opening (panel B-C).

**Figure 9.**
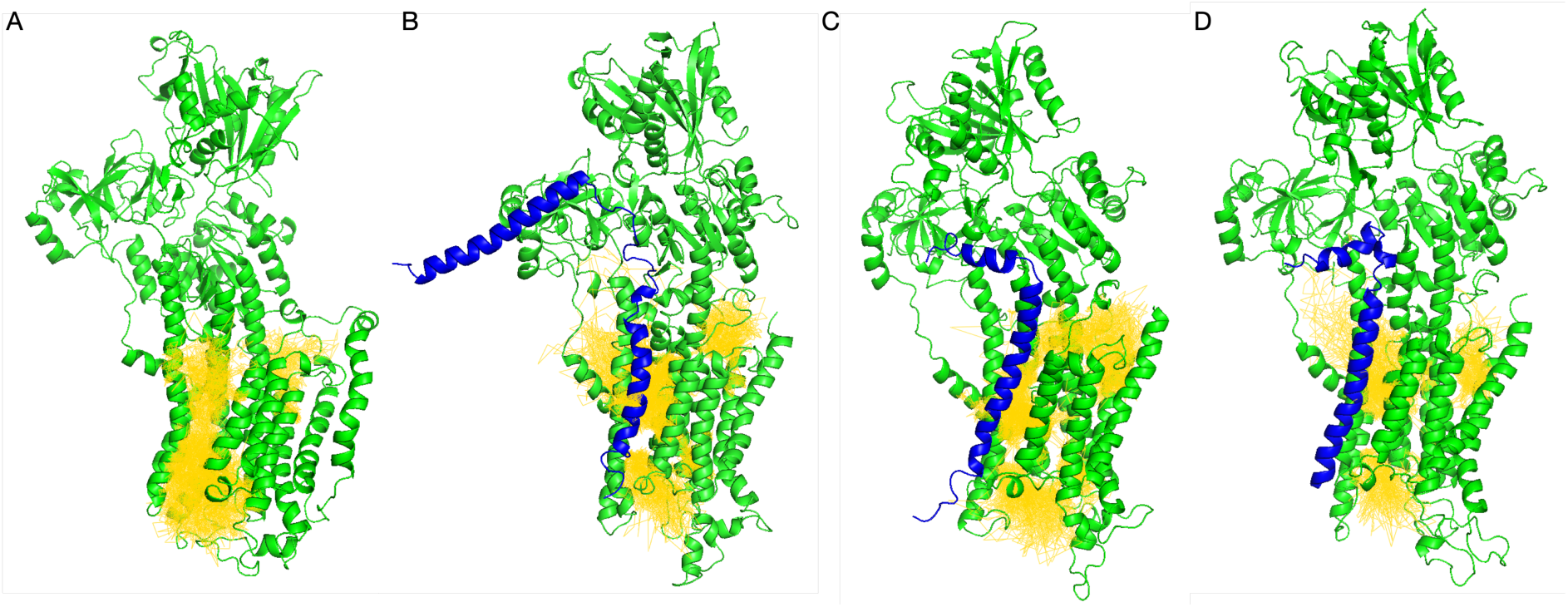
Representation of the water fluxes during MD simulations of the different systems. A) Apo-SERCA2b, B) SERCA2b∷NNAT α, C) SERCA2b∷NNAT β D) SERCA2b∷PLB. It can be observed how there is no connection between the N- and the C-paths in the apo-SERCA2b system, while they are fully connected on all three other systems, especially when in complex with PLB, where they lose all contacts with the endoplasmic exit. Green: SERCA2b; blue: NNAT/PLB; yellow: water molecule paths.

A salt bridge was formed for most of the MD simulation between Glu28 of NNAT α and the residues Arg324 and Lys328 of SERCA (Fig. S4), known to be important for conformational rearrangements of the M4S4 domain. On the other hand, most of the interactions formed between the β isoform and SERCA are hydrophobic interactions produced by residues present in the transmembrane helices.

Overall, these results support the hypothesis that the two NNAT isoforms behave similarly to PLB, leading to a decrease in the rate of Ca^2+^ transport across the membrane by causing a specific conformational rearrangement affecting the “N” and “C” pathways.

## Conclusions

In this study, we aimed to elucidate the structural and functional characteristics of the two NNAT isoforms and their interaction with the SERCA2b Ca^2+^ pump. The RosettaFold-generated models, which closely resembled the experimentally solved regulin structures, like PLB and sarcolipin, were used as starting point for MD simulations for refinement. The analysis of the simulation showcased stable conformations, after an initial structural rearrangement, and confirmed a predominant α-helical secondary structure.

A study of the aggregation potential of both NNAT isoforms using both all-atoms and coarse grained MD simulations revealed a notably fast aggregation process. These results support experimental evidence linking NNAT aggregates to degenerative diseases such as Lafora disease, highlighting coarse-grained simulations as a viable method to assess protein aggregation.

To investigate the interaction between NNAT and SERCA2b, we carried out protein∷protein docking using the transmembrane helix of NNAT. The results identified a NNAT binding site within the groove between helices M2, M6, and M9 of SERCA2b, a region known for PLB binding. This supports the hypothesis that NNAT isoforms interact with SERCA2b in a manner similar to regulins, potentially modulating the pump activity by altering its conformational dynamics.

Further MD simulations of the NNAT-SERCA2b complexes revealed that both NNAT isoforms influence water flux through the pump, akin to the inhibitory effect observed with PLB. The α isoform showed a slightly greater impact on water molecule retention in the cytoplasm. Additionally, the formation of a salt bridge between Glu28 of NNAT α and key residues in SERCA2b (Arg324 and Lys328) suggests specific molecular interactions that could underlie the regulatory effect.

In conclusion, our comprehensive computational findings point out that NNAT isoforms modulate SERCA2b activity via structural interactions similar to those of PLB, suggesting that neuronatin could be considered a “regulin-like” protein, if not even a new member of the regulin family. This study widens our understanding of the functional role of NNAT in Ca^2+^ homeostasis and provides foundations for future experimental investigations to explore these regulatory mechanisms and design small molecules affecting NNAT activity.

## Authors contributions

OBM: Computational simulations, data processing, writing – original draft preparation. LC: Data visualization, formal analysis, writing – original draft preparation. EDF: Study conceptualization, writing – review and editing. MC: Study conceptualization, writing – review and editing. IE: Project oversight, study conceptualization, writing – review and editing.

