## Supplemental Information for "Identification of neuronatin as a SERCA2b regulin-like protein and assessment of its aggregation propensity via coarse grained simulations"

### Supplementary tables

| Pose number | PIPER cluster size |
|---|---|
| 1 | 256 |
| 2 | 222 |
| 3 | 141 |
| 4 | 134 |
| 5 | 97 |
| 6 | 86 |
| 7 | 27 |
| 8 | 18 |
| 9 | 12 |
| 10 | 6 |

Supplementary table 1. PIPER protein::protein docking poses sorted by cluster size.

### Supplementary figures

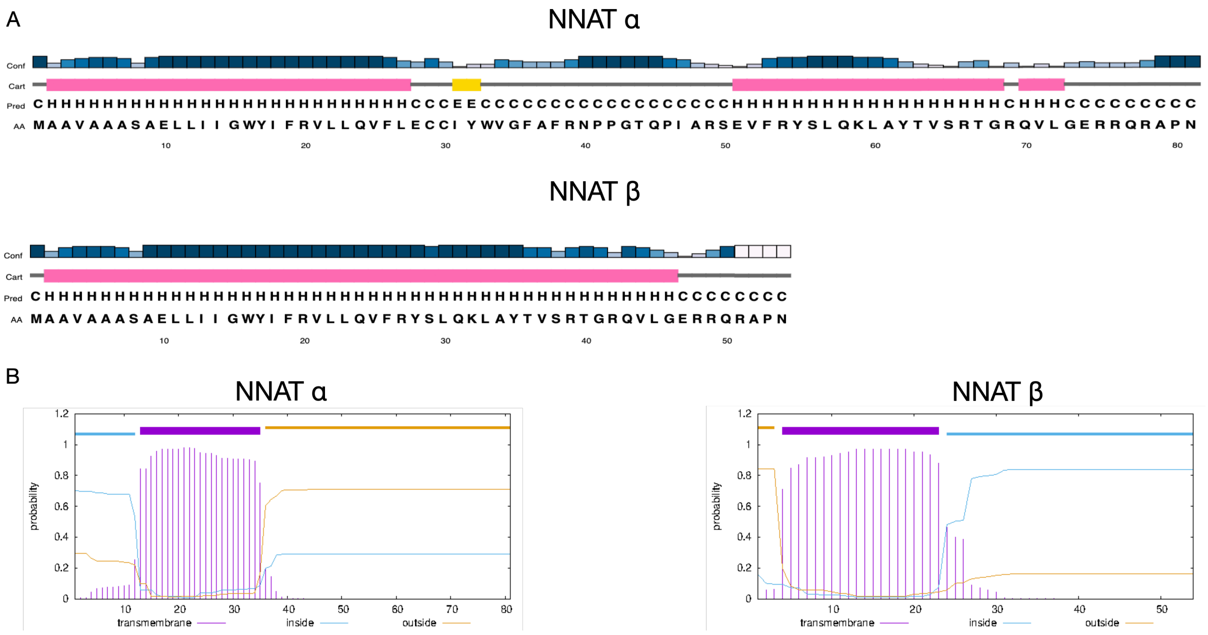


Supplementary figure 1. Secondary structure prediction by PsiPred (A) and transmembrane helix prediction by TMHMM (B) for the two NNAT isoforms. All predictions, with small differences, predict the presence of a transmembrane loop at the N-terminal. In the longer α isoform, the two helices are separated by a long unfolded loop, while in the β isoform, no such loop is predicted.

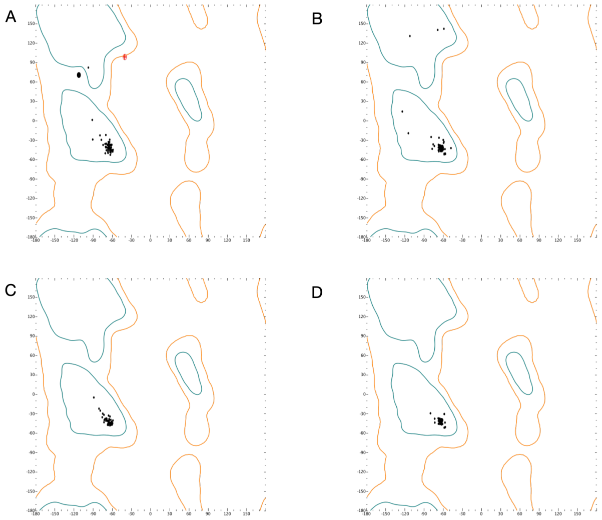


Supplementary figure 2. Ramachandran plot for the AI-generated models. A) NNAT α generated by AlphaFold. One outlier (Arg39) can be observed. B) NNAT α generated by RoseTTAFold. C) NNAT β generated by AlphaFold. D) NNAT β generated by RoseTTAFold

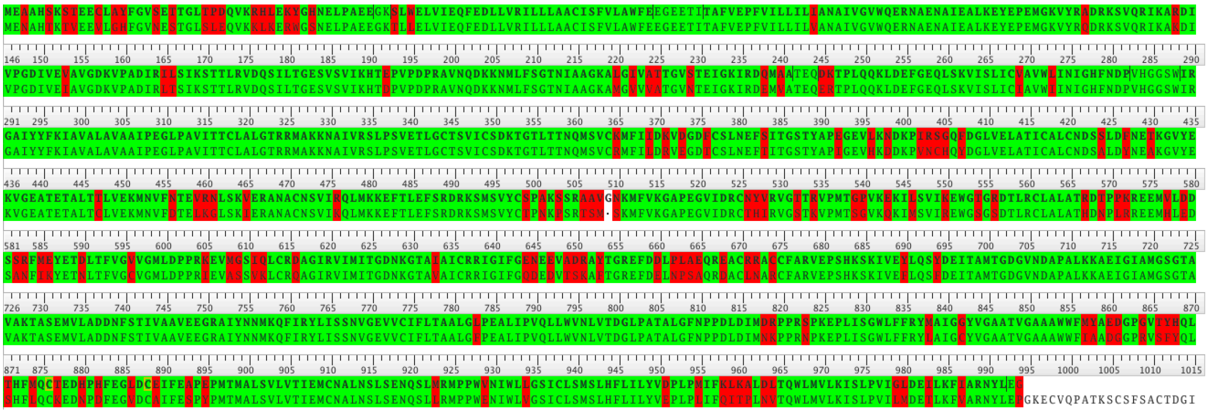

Supplementary figure 3. Sequence alignment between the sequence of *O. cuniculus* SERCA1a as present in the crystal structure with PDB code 4KYT and human SERCA2b. The identical residues are colored in greed, while the non-identical in red. Even if not identical, most residues are highly conserved (e.g., I→L, D→E, ...). ID%=80.2%, similarity % = 89.2%.

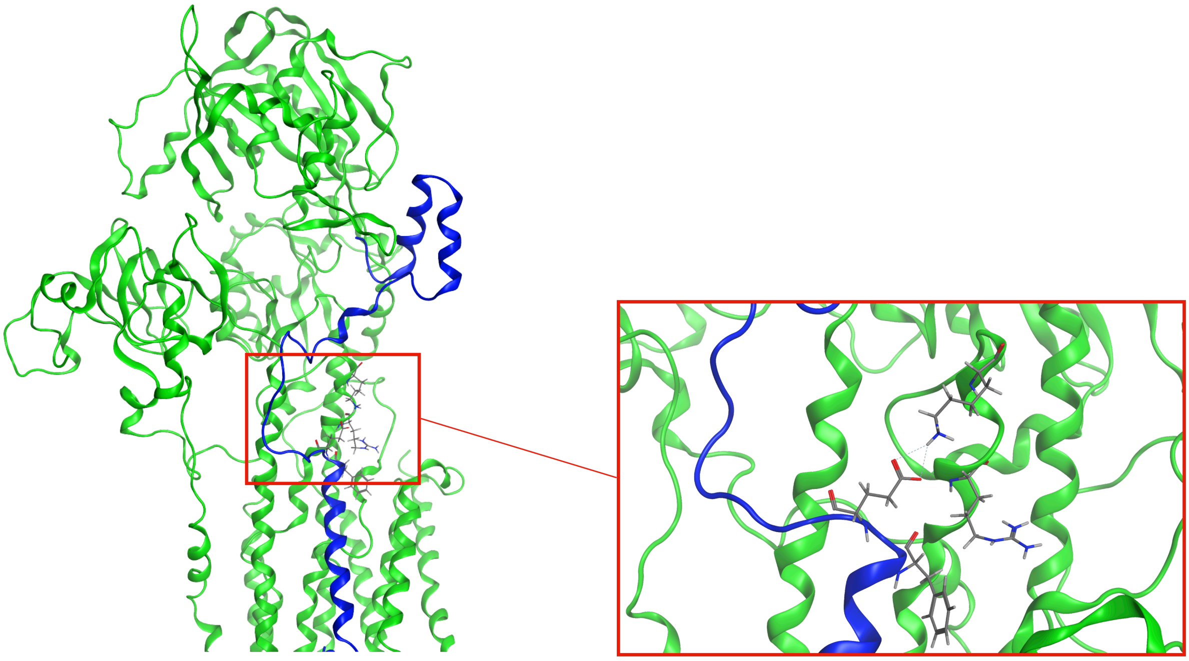

Supplementary figure 4. Salt bridge interaction between Glu28 of α NNAT (blue) and Arg324 and Lys328 of SERCA (green). These residues have been described in literature as key residues for SERCA conformational changes.
